# Hepatocyte-specific deletion of XBP1 sensitizes mice to liver injury through hyperactivation of IRE1α

**DOI:** 10.1101/2020.04.23.058446

**Authors:** Caroline C. Duwaerts, Kevin Siao, Russell K. Soon, Chris Her, Takao Iwawaki, Kenji Kohno, Aras N. Mattis, Jacquelyn J. Maher

## Abstract

X-box binding protein-1 (XBP1) is a transcription factor that plays a central role in controlling cellular responses to endoplasmic reticulum (ER) stress. Under stress conditions, the transcriptionally active form of XBP1 is generated by unique splicing of *Xbp1* mRNA by the ER-resident protein inositol-requiring enzyme-1 (IRE1α). Genetic deletion of XBP1 has multiple consequences: some resulting from the loss of the transcription factor per se, and others related to compensatory upstream activation of IRE1α. The objective of the current study was to investigate the effects of XBP1 deletion in adult mouse liver and determine to what extent they are direct or indirect. XBP1 was deleted specifically from hepatocytes in adult *Xbp1*^fl/fl^ mice using AAV8-Transthyretin-Cre (*Xbp1*^Δhep^). *Xbp1*^Δhep^ mice exhibited no liver disease at baseline, but developed acute biochemical and histologic liver injury in response to a dietary challenge with fructose for 4 wk. Fructose-mediated liver injury in *Xbp1*^Δhep^ mice coincided with heightened IRE1α activity, as demonstrated by cJun phosphorylation and regulated IRE1α -dependent RNA decay (RIDD). Activation of eIF2α was also evident, with associated up-regulation of the pro-apoptotic molecules CHOP, BIM and PUMA. To determine whether the adverse consequences of liver-specific XBP1 deletion were due to XBP1 loss or heightened IRE1α activity, we repeated a fructose challenge in mice with liver-specific deletion of both XBP1 and IRE1α (*Xbp1*^Δhep^;*IRE1α*^Δhep^). *Xbp1*^Δhep^;*IRE1α*^Δhep^ mice were protected from fructose-mediated liver injury and failed to exhibit any of the signs of ER stress seen in mice lacking XBP1 alone. The protective effect of IRE1α deletion persisted even with long-term exposure to fructose. *Xbp1*^Δhep^ mice developed liver fibrosis at 16 wk, but *Xbp1*^Δhep^;*IRE1α*^Δhep^ mice did not. Overall, the results indicate that the deleterious effects of hepatocyte-specific XBP1 deletion are due primarily to hyperactivation of IRE1α. They support further exploration of IRE1α as a contributor to acute and chronic liver diseases.

## Introduction

X-box binding protein-1 (XBP1) is an important component of the signal transduction network that protects cells against ER stress. XBP1 is positioned downstream of IRE1α (inositol-requiring enzyme-1), one of three canonical ER stress sensors (IRE1, ATF6, PERK) residing in the ER membrane. IRE1α has kinase and endoribonuclease activities that are unleashed under conditions of ER stress. When IRE1α is activated, its endoribonuclease acts upon *Xbp1* mRNA by splicing a 26-nucleotide fragment that enables translation of a protein termed XBP1s. XBP1s is a transcription factor that induces genes involved in chaperoning proteins through the ER and degrading proteins that cannot be properly folded in the ER [1]. XBP1s also induces genes pertinent to phospholipid synthesis, which enhance ER membrane biogenesis and increase the capacity of the organelle [2,3].

Cell-specific deletion of XBP1 often results in adverse consequences. Deletion of XBP1 from lymphoid precursors prevents the maturation of B cells into plasma cells [4]; deletion from intestinal epithelia predisposes to inflammatory bowel disease [5]; and deletion of XBP1 from CNS neurons promotes leptin resistance and obesity [6]. In these situations, the targeted loss of XBP1 causes enhanced ER stress in the affected cells, which can lead to cell death and associated inflammation. Targeted disruption of XBP1, however, can in some cases lead to mixed positive and negative outcomes. This is true of the liver, in which XBP1 deletion from hepatocytes improves hepatic insulin sensitivity [7] and reduces the hepatic contribution to circulating lipids [8] but sensitizes the liver to pharmacologic ER stress [9] and impairs liver regeneration [10].

The pleiotropic consequences of XBP1 deletion in the liver may be due in part to the large number of genes targeted by XBP1 and the impact of these genes on diverse biological processes [10,11]. There is also the possibility that XBP1 deletion affects the liver indirectly through compensatory effects on other molecules involved in the ER stress response. Indeed, extensive crosstalk occurs among the signaling events triggered by the three canonical ER stress transducers IRE1, ATF6 and PERK [12]. Importantly, disruption of one arm of the network can lead to exaggerated activity of the others, which can result in a maladaptive or even fatal response to stress [13]. With this in mind, one important consequence of XBP1 deletion is hyperactivation of IRE1α [8]. Hyperactivation of IRE1α leads to significant broadening of its endoribonuclease activity, which in turn prompts large-scale degradation of mRNAs in a process called regulated IRE1α-dependent decay (RIDD) [14]. IRE1α hyperactivation also accentuates its kinase activity toward TRAF2, initiating a cascade of events culminating in the activation of JNK and downstream targets such as cJun [15]. Both of these events can trigger cell death.

Experts reporting that XBP1 deletion is beneficial to the liver have attributed the effect to a reduction in hepatic lipogenesis in the absence of XBP1 [8,11]. Diminishing XBP1 activity may reduce lipogenesis under mild stress conditions, but it is uncertain whether this metabolic improvement is offset by other negative consequences. The goal of the current study was to examine the impact of hepatocyte-specific XBP1 deletion in the adult mouse liver, under basal conditions and in response to a mild metabolic stress (fructose feeding). We wished to gain insight into metabolic outcomes as well as cell survival, and to dissect whether the phenotype of XBP1-deficient mice was due primarily to XBP1 loss or to upregulation of other ER stress pathways.

## Methods

### Mice and experimental diets

XBP1 conditional knockout mice on a C57BL/6 background (*Xbp1*^fl/fl^) were obtained from Drs. Ann-Hwee Lee and Laurie Glimcher [8]. IRE1α conditional knockout mice (*IRE1α*^fl/fl^) were generated as previously described [16] and back-crossed for 10 generations to C57BL/6. The two strains were cross-bred to generate *Xbp1*^fl/fl^;*IRE1α*^fl/fl^ conditional knockout mice. At 8 wk of age, *Xbp1*^fl/fl^ or *Xbp1*^fl/fl^;*IRE1α*^fl/fl^ mice were injected IV with either 4 x 10^11^ GC AAV8-Transthyretin-Cre or 4 x 10^11^ GC AAV8-CMV-null as a control (Vector Biolabs, Malvern, PA). Gene-deleted mice are designated *Xbp1*^Δhep^ and *Xbp1*^Δhep^;*IRE1α*^Δhep^. Animals were housed for 2 wk after AAV8 treatment before initiating experimental studies. At 10 wk of age, mice were placed on either a chow diet (Pico Lab Diets #5053) or a fructose-enriched diet (Envigo TD.89247) for intervals up to 16 wk. At the end of each experiment, mice were fasted for 4 h before killing. Positive controls for ER stress were generated by injecting adult C57BL/6 mice with tunicamycin (1 mg/kg) and killing them 3 h later. All mouse experiments were performed in accordance with guidelines set by the America Veterinary Medical Association. All mouse studies were reviewed and approved by the Committee on Animal Research at the University of California San Francisco.

### Gene expression

RNA was extracted from whole liver in TRIzol (Invitrogen, Carlsbad, CA). RNA was then purified using a Direct-zol RNA Miniprep kit (ZymoResearch, Irvine, CA) and cDNA synthesized as previously described [17]. Gene expression was assessed by quantitative PCR using PrimeTime qPCR assays (Integrated DNA Technologies, Coralville, IA), E@sy Oligo primers (Millipore-Sigma, Burlington, MA) or TaqMan Assays (Life Technology, Carlsbad, CA), followed by normalization to mouse β-glucuronidase.

### Histology and immunohistochemistry

Formalin-fixed sections of liver tissue were stained with hematoxylin and eosin. Cell death was evaluated by terminal deoxynucleotidyl transferase dUTP nick end labeling (TUNEL) (ApopTag Plus Peroxidase *In Situ* Apoptosis Detection Kit, Millipore-Sigma). Cell proliferation was evaluated by Ki-67 immunostaining (Cell Signaling Technology, Danvers, MA). Sections were photographed using a Nikon Microphot microscope (Nikon, Melville, NY) equipped with a SPOT digital camera (Diagnostic Instruments, Inc., Sterling Heights, MI). TUNEL-positive and Ki-67-positive cells were counted manually in 10 microscopic fields per liver, each measuring 0.4 mm^2^. Data were reported as the average number of cells per microscopic field.

### Quantitation of hepatic lipids

Lipids were extracted from fresh liver tissue using the Folch method [18]. Total triglyceride was measured spectrophotometrically as previously described (TR0100; Millipore-Sigma) [19].

### Quantitation of hepatic fibrosis

Hepatic fibrosis was assessed morphometrically in Sirius Red-stained tissue sections using LAS X (Leica Microsystems, Wetzlar, Germany). The fibrosis area (%) for each liver was assessed as the mean measurement of 6 microscopic fields, each measuring 0.4 mm^2^. Fibrosis was also quantitated by measuring the amount of hydroxyproline in tissue homogenates [20]. Values are reported as mg hydroxyproline/g liver.

### Serum tests

Alanine aminotransferase (ALT), total cholesterol and total triglycerides were measured in mouse serum using an ADVIA 1800 autoanalyzer (Siemens Healthcare Diagnostics, Deerfield, IL) in the clinical chemistry laboratory at the Zuckerberg San Francisco General Hospital.

### Western blotting

Livers were homogenized in RIPA buffer containing protease and phosphatase inhibitors. Aliquots were separated by electrophoresis (Bio-Rad TGX, Hercules, CA) and transferred to PVDF membranes (Bio-Rad). Proteins were identified using the following primary antibodies (Cell Signaling Technology, Danvers, MA and Santa Cruz Biotechnology, Dallas, TX): activating transcription factor 3 (ATF3), activating transcription factor 4 (ATF4), the BH3-only proteins BIM and Bcl-XL, C/EBP homologous protein (CHOP), eukaryotic translation initiating factor 2α (eIF2α), P-eIF2α, cJun, P-cJun, IRE1α, P-IRE1α, lamin B1, p53 upregulated modulator of apoptosis (PUMA), tubulin and XBP1. Peroxidase-conjugated secondary antibodies were from Cell Signaling Technology. Antigen-antibody complexes were visualized by chemiluminescence using a FluorChem FC2 system (Protein Simple, San Jose, CA) and Super Signal West Dura (Thermo Scientific).

### Statistical Analysis

All experimental results were compared using 1-way analysis of variance (ANOVA) followed by Tukey’s multiple comparisons test unless otherwise stated. Statistical analyses were performed with Prism 8.3.0 software (GraphPad Software, San Diego, CA).

## Results

AAV8-Ttr-Cre successfully deleted XBP1 from hepatocytes in *Xbp1*^fl/fl^ mice, as shown by the near-complete absence of nuclear XBP1s in the livers of *Xbp1*^Δhep^ mice exposed to inducers of ER stress (**Figure S1A, B**). *Xbp1*^Δhep^ mice had low levels of serum lipids (**Figure S1C**), which has been reported previously and attributed to impaired hepatic lipid secretion [8,11,21]. Liver histology in *Xbp1*^Δhep^ mice was normal (**Figure S1D**). When *Xbp1*^Δhep^ mice were fed a fructose-enriched diet for 1 wk, lipogenic genes were induced in the liver, although to a lesser extent than *Xbp1*^fl/fl^ controls (**Figure 1A**). Serum cholesterol and hepatic triglyceride levels increased modestly in both groups of mice in response to fructose feeding, but serum triglycerides were unchanged and liver histology and ALT remained normal (**Figure 1B, C**).

**Figure 1.**
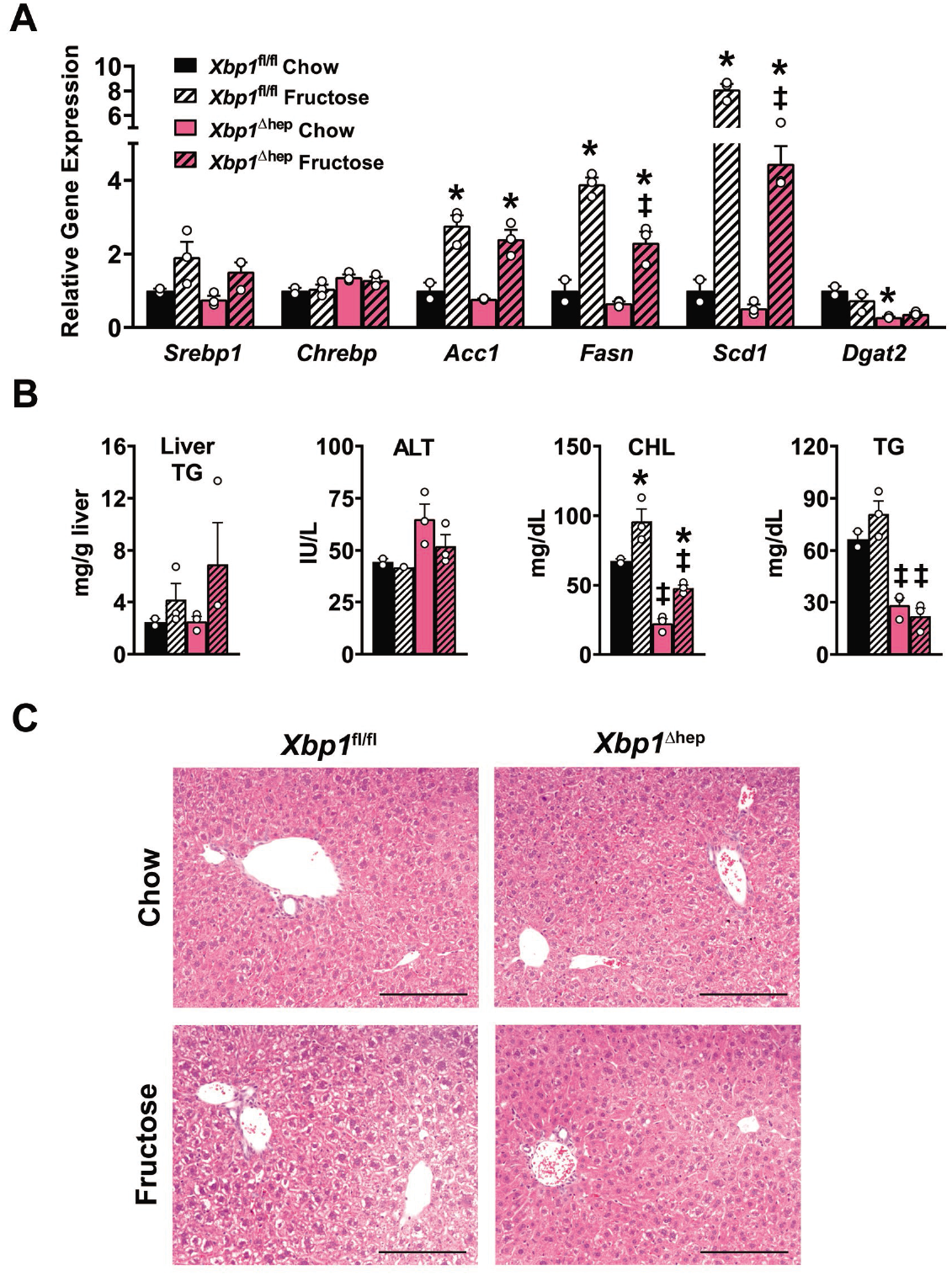
Response of *Xbp1*^Δhep^ mice to fructose feeding for 1 week. **(A)** Histogram demonstrates lipogenic gene expression in *Xbp1*^fl/fl^ and *Xbp1*^Δhep^ livers after 1 wk of fructose feeding. **(B)** Graphs depict the corresponding hepatic triglyceride levels after fructose feeding, as well as serum levels of cholesterol (CHL), triglyceride (TG) and ALT. **(C)** Liver histology following 1 wk of fructose feeding. Bar = 200 μm. Values represent mean ± SEM for n = 3. *P* < 0.001 by one-way ANOVA for *Acc1, Fasn, Scd1*, CHL and TG. Using Tukey’s multiple comparisons test, * *P* < 0.05 for fructose vs. chow of same genotype and ‡ *P* < 0.05 for *Xbp1*^Δhep^ vs. *Xbp1*^fl/fl^.

When fructose feeding was continued for 4 wk, lipogenic gene expression remained elevated in both groups of mice but mRNA levels continued to be higher in *Xbp1*^fl/fl^ than *Xbp1*^Δhep^ livers (**Figure S2**). Despite their weaker lipogenic response to fructose, *Xbp1*^Δhep^ mice displayed overt hepatic steatosis at the 4-wk time point (**Figure 2A, B**). Also evident in *Xbp1*^Δhep^ livers were cell death and regeneration, documented by TUNEL staining and Ki67 immunohistochemistry. Serum ALT levels rose to three times normal in fructose-fed *Xbp1*^Δhep^ mice at 4 wk. In contrast, ALT and liver histology remained normal in fructose-fed *Xbp1*^fl/fl^ mice, although hepatic triglyceride levels were elevated above the chow-fed baseline (**Figure 2A, B**).

**Figure 2.**
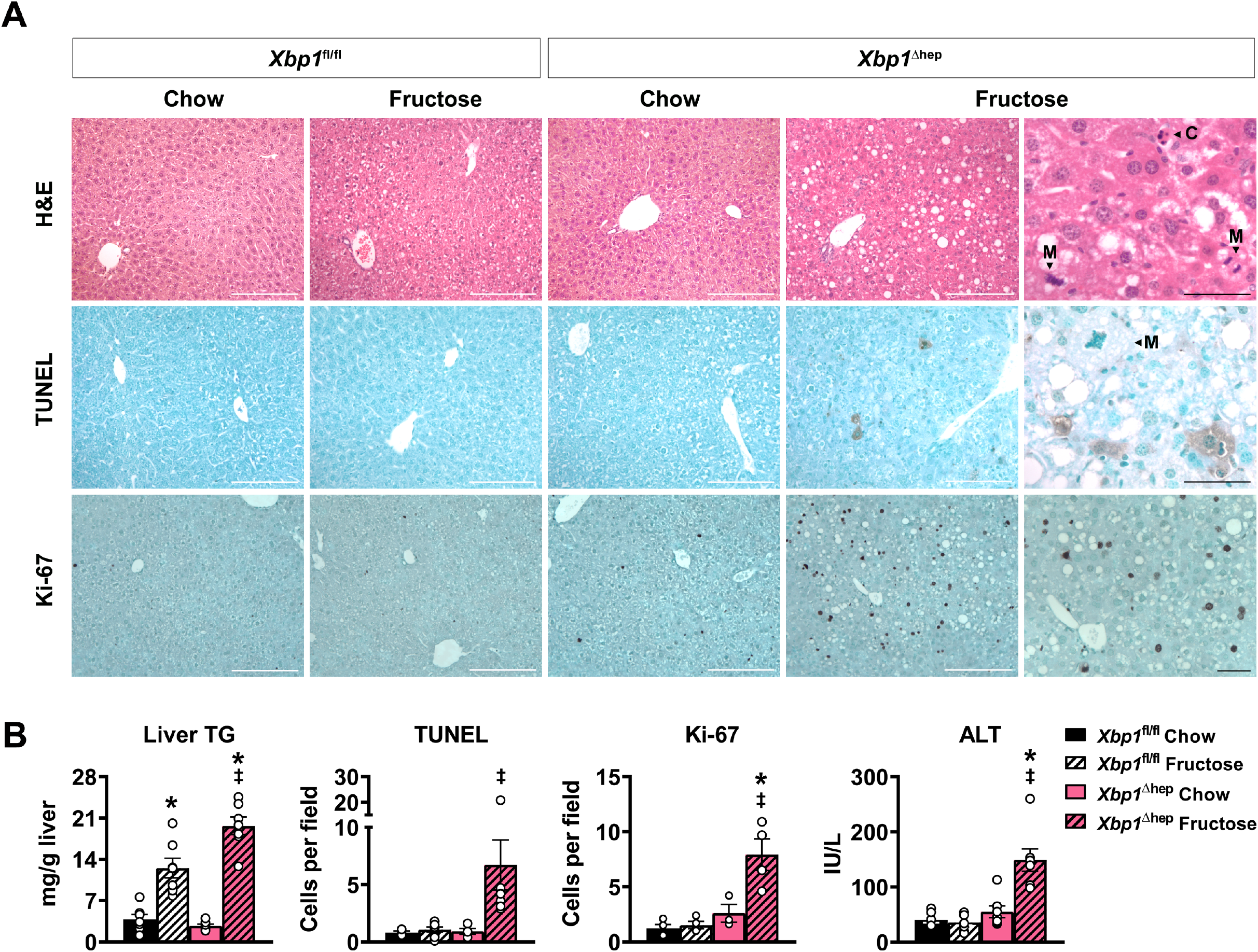
Response of *Xbp1*^Δhep^ mice to fructose feeding for 4 weeks. **(A)** Photomicrographs illustrate H&E-stained liver sections as well as TUNEL and Ki-67 immunohistochemistry in *Xbp1*^fl/fl^ and *Xbp1*^Δhep^ livers after 4 wk of fructose feeding. White bars = 200 μm. Higher-power views illustrate cell death (C) and mitoses (M) and highlight Ki-67-positive cells. Black bars = 50 μm. **(B)** Graphs depict quantitative measures of hepatic triglyceride, TUNEL and Ki-67 staining as well as serum ALT levels. Values represent mean ± SEM for n = 7. *P* < 0.05 by one-way ANOVA for liver TG, TUNEL, Ki67 and ALT. Using Tukey’s multiple comparisons test, * *P* < 0.05 for fructose vs. chow of same genotype and ‡ *P* < 0.05 for *Xbp1*^Δhep^ vs. *Xbp1*^fl/fl^.

To explore the connection between hepatocyte XBP1 deletion and the development of liver injury in fructose-fed *Xbp1*^Δhep^ mice, we investigated the influence of fructose feeding on the IRE1α-XBP1 axis and its downstream target JNK. In *Xbp1*^fl/fl^ mice, fructose feeding stimulated nuclear translocation of spliced XBP1 in the liver at 1 wk but not 4 wk (**Figure 3A**). Fructose feeding also transiently increased hepatic expression of IRE1α and induced phosphorylation of the JNK target cJun at 1 wk but not 4 wk. In *Xbp1*^Δhep^ mice, spliced XBP1 was undetectable in response to fructose feeding at any time point. IRE1α expression, however, was markedly elevated in the livers of *Xbp1*^Δhep^ mice, independent of diet or time interval. *Xbp1*^Δhep^ mice were confirmed to have impaired XBP1 transcriptional activity based on the significant suppression of direct XBP1 target genes in the liver compared to *Xbp1*^fl/fl^ mice (**Figure 3B**). At the same time, IRE1α hyperactivation was demonstrable in *Xbp1*^Δhep^ livers by the significant downregulation of genes known to be RIDD targets (**Figure 3B**). When *Xbp1*^Δhep^ mice were fed chow, IRE1α induction coincided with only mild and variable activation of cJun over 4 wk. When *Xbp1*^Δhep^ mice were fed fructose, however, cJun was definitively activated at 4 wk, in parallel with the development of liver injury.

**Figure 3.**
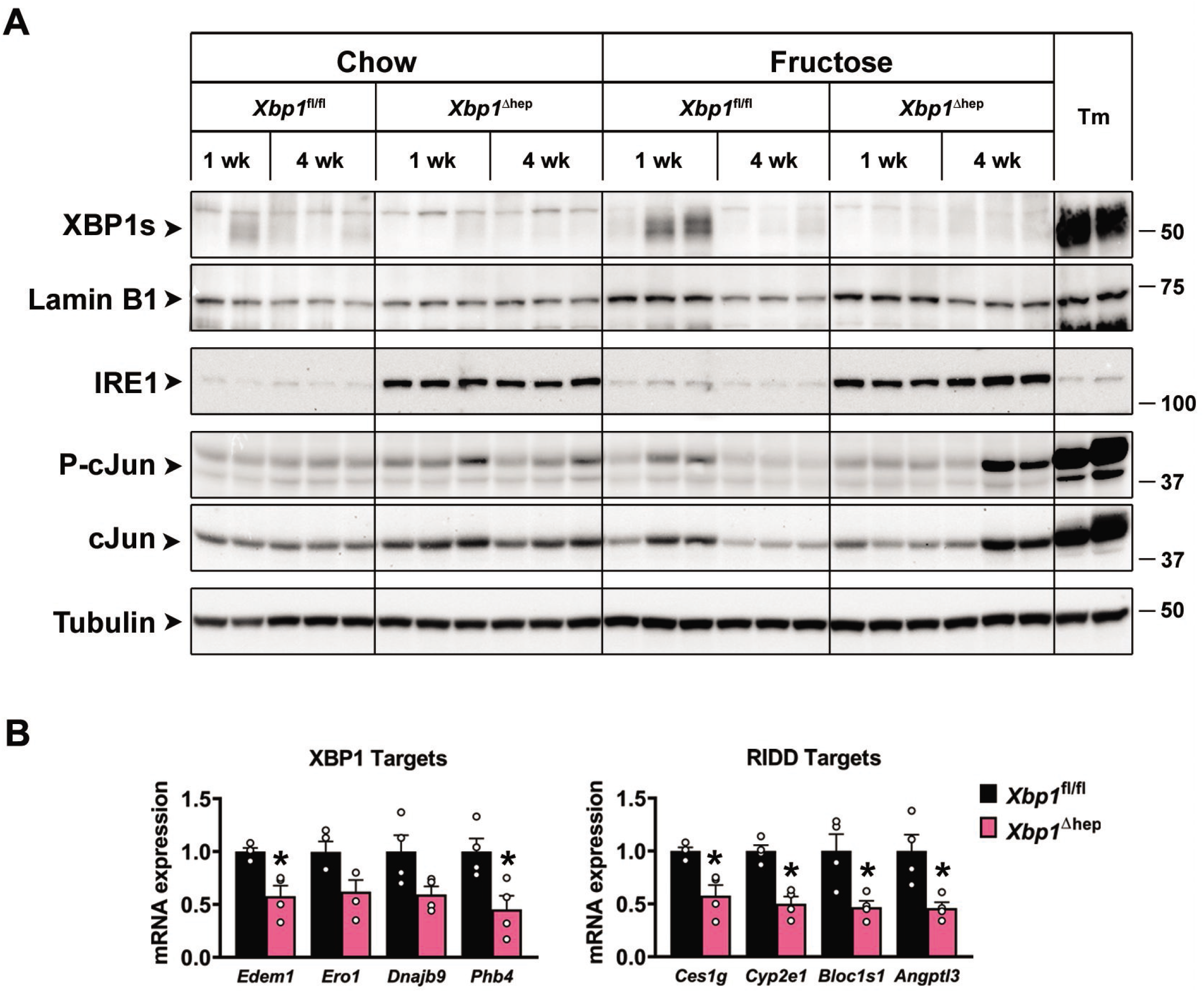
Effect of XBP1 deletion and fructose feeding on the IRE1α-XBP1 axis. **(A)** Western blots illustrate the effects XBP1 deletion, with or without fructose feeding for 1 or 4 wk, on activity of the IRE1α-XPB1 axis. Tunicamycin treatment (Tm) served as a positive control. XBP1s and lamin B1 were measured in nuclear extracts; other proteins were measured in whole liver homogenates. In *Xbp1*^fl/fl^ mice, fructose feeding stimulated nuclear translocation of XBP1s at 1 wk but not 4 wk. This coincided with modest but and transient up-regulation of IRE1α and phosphorylation of cJun. In *Xbp1*^Δhep^ mice, was strongly upregulated regardless of diet or duration. This resulted in robust activation of cJun, but only after 4 wk of fructose feeding. **(B)** Graphs depict relative mRNA expression for a panel of direct XBP1 target genes and separate panel of RIDD target genes in the livers of *Xbp1*^fl/fl^ and *Xbp1*^Δhep^ mice. Values represent mean ± SEM for n = 4. By unpaired t-test, * *P* < 0.05 for *Xbp1*^Δhep^ vs. *Xbp1*^fl/fl^.

Because cell death is not exclusively linked to IRE1α in the setting of ER stress [15,22], we also investigated the influence of fructose feeding on other ER stress pathways in *Xbp1*^Δhep^ mice. We focused on eIF2α, whose activation can lead to cell death by up-regulating apoptotic proteins and downregulating survival proteins. We observed that XBP1 deletion by itself promoted phosphorylation of eIF2α even with chow feeding, as has been reported previously [23]. In chow-fed *Xbp1*^Δhep^ mice, however, eIF2α phosphorylation did not trigger any downstream events such as up-regulation of ATF 3/4 or CHOP (**Figure 4A**). When XBP1 deletion was coupled with fructose feeding, eIF2α phosphorylation was accompanied by the induction of ATF3 and ATF4 and up-regulation of the apoptotic proteins CHOP, BIM and PUMA, without any apparent change in the anti-apoptotic protein Bcl-XL. These changes were accompanied by a significant increase in hepatic mRNA encoding the pro-inflammatory protein CCL2 and smaller increases in other inflammatory and fibrotic genes (**Figure 4B**). Taken together these findings indicate that hepatocyte-specific XBP1 deletion is sufficient to induce activation of IRE1α and eIF2α, but these alterations are without consequence until an independent stimulus such as fructose feeding is applied. In the presence of fructose, XBP1 deletion leads to hepatic steatosis, liver cell death and up-regulation of inflammatory signals. Notably, fructose feeding *per se* is a mild metabolic challenge, insufficient to provoke a robust or sustained ER stress response in control mice. Hepatocyte-specific XBP1 deletion, therefore, sensitizes the liver to what otherwise would be an innocuous metabolic insult.

**Figure 4.**
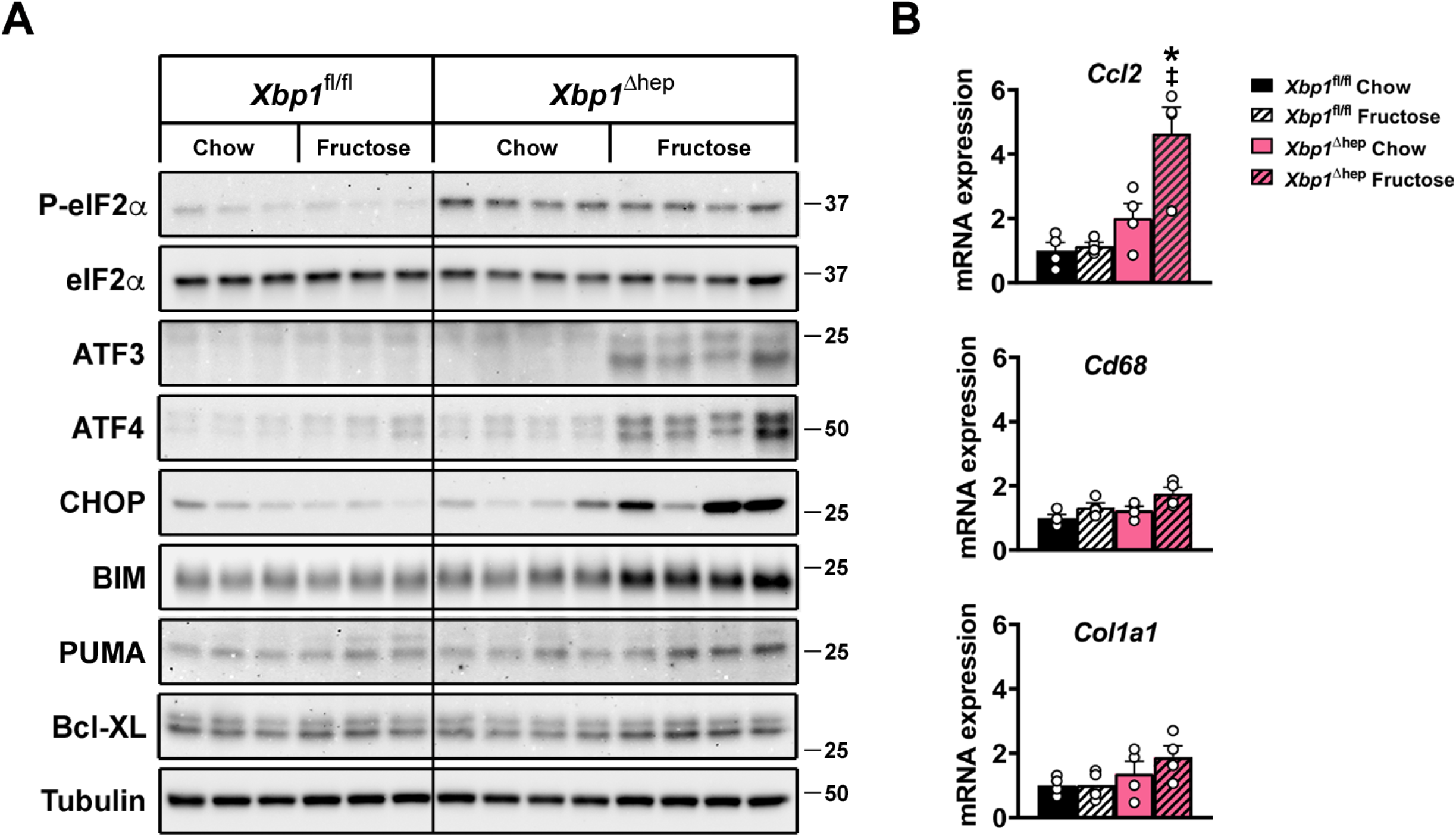
Effect of XBP1 deletion and fructose feeding on other ER stress pathways. **(A)** Western blots illustrate the effects XBP1 deletion, with or without fructose feeding for 4 wk, on the activity of eIF2α and the expression of several eIF2α targets. *Xbp1*^Δhep^ mice exhibited phosphorylation of eIF2α with chow feeding as well as fructose feeding at 4 wk. Up-regulation of molecules downstream of eIF2α occurred only after fructose feeding in *Xbp1*^Δhep^ mice. **(B)** Graphs demonstrate the effect of fructose feeding for 4 wk on select genes pertinent to hepatic inflammation and fibrosis. *Ccl2* mRNA is significantly induced by fructose feeding in *Xbp1*^Δhep^ mice, but there is no significant change in the expression of *Cd68* or *Col1a1*. Values represent mean ± SEM for n = 4. *P* < 0.001 by one-way ANOVA for *Ccl2*. * *P* < 0.05 for fructose vs. chow of same genotype and ‡ *P* < 0.05 for *Xbp1*^Δhep^ vs. *Xbp1*^fl/fl^.

Because up-regulation of IRE1α was so pronounced in the livers of *Xbp1*^Δhep^ mice, we reasoned that IRE1α activation was central to their sensitization to liver injury. To test this directly, we generated mice with AAV8-mediated deletion of XBP1 as well as the RNase domain of IRE1α (*Xbp1*^Δhep^;*IRE1α*^Δhep^) [16]. *Xbp1*^Δhep^;*IRE1α*^Δhep^ mice expressed a truncated form of IRE1α indicative of the RNase domain deletion [24] (**Figure S3A**). In contrast to *Xbp1*^Δhep^ mice, *Xbp1*^Δhep^;*IRE1α*^Δhep^ mice exhibited no spontaneous XBP1 mRNA splicing (**Figure S3B**). Direct XBP1 target genes were suppressed in the livers of *Xbp1*^Δhep^;*IRE1α*^Δhep^ mice, as expected due to the absence of XBP1; RIDD target genes were not suppressed, indicating that RIDD was inactive in the doubly-deficient mice (**Figure S3C**). When *Xbp1*^Δhep^;*IRE1α*^Δhep^ mice were challenged with a fructose-enriched diet for 4 wk, they were completely spared from fructose-induced hepatic steatosis and liver injury (**Figure 5A, B**). *Xbp1*^Δhep^;*IRE1α*^Δhep^ mice did not exhibit any of the signs of ER stress noted in *Xbp1*^Δhep^ mice. Nor did they display any significant induction of select inflammatory or fibrotic genes (**Figure 6A, B**).

**Figure 5.**
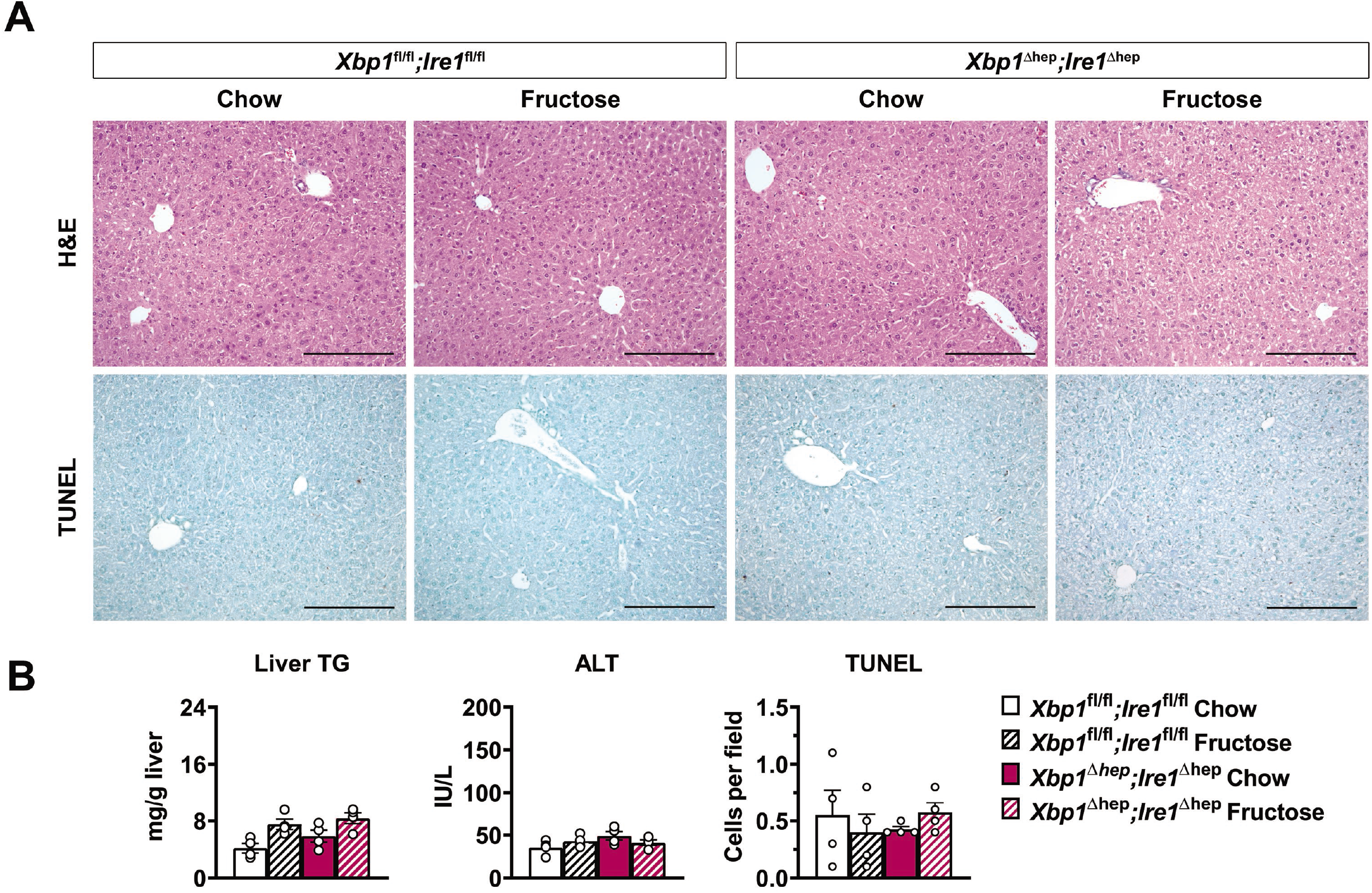
Fructose feeding does not induce liver injury in mice with dual deletion of XBP1 and IRE1α. **(A)** Photomicrographs depict liver histology in *Xbp1*^fl/fl^;*IRE1α*^fl/fl^ and *Xbp1*^Δhep^;*IRE1α*^Δhep^ mice after 4 wk of chow or fructose feeding. There is no evidence of hepatic steatosis by histology or quantitative triglyceride, and no evidence of liver cell death, as assessed by TUNEL staining. Bar = 200 μm. **(B)** Measurements of liver TG, serum ALT and TUNEL-positive cell counts confirm the absence of hepatic steatosis or liver injury in *Xbp1*^Δhep^;*IRE1α*^Δhep^ mice. Values represent mean ± SEM for n = 4.

**Figure 6.**
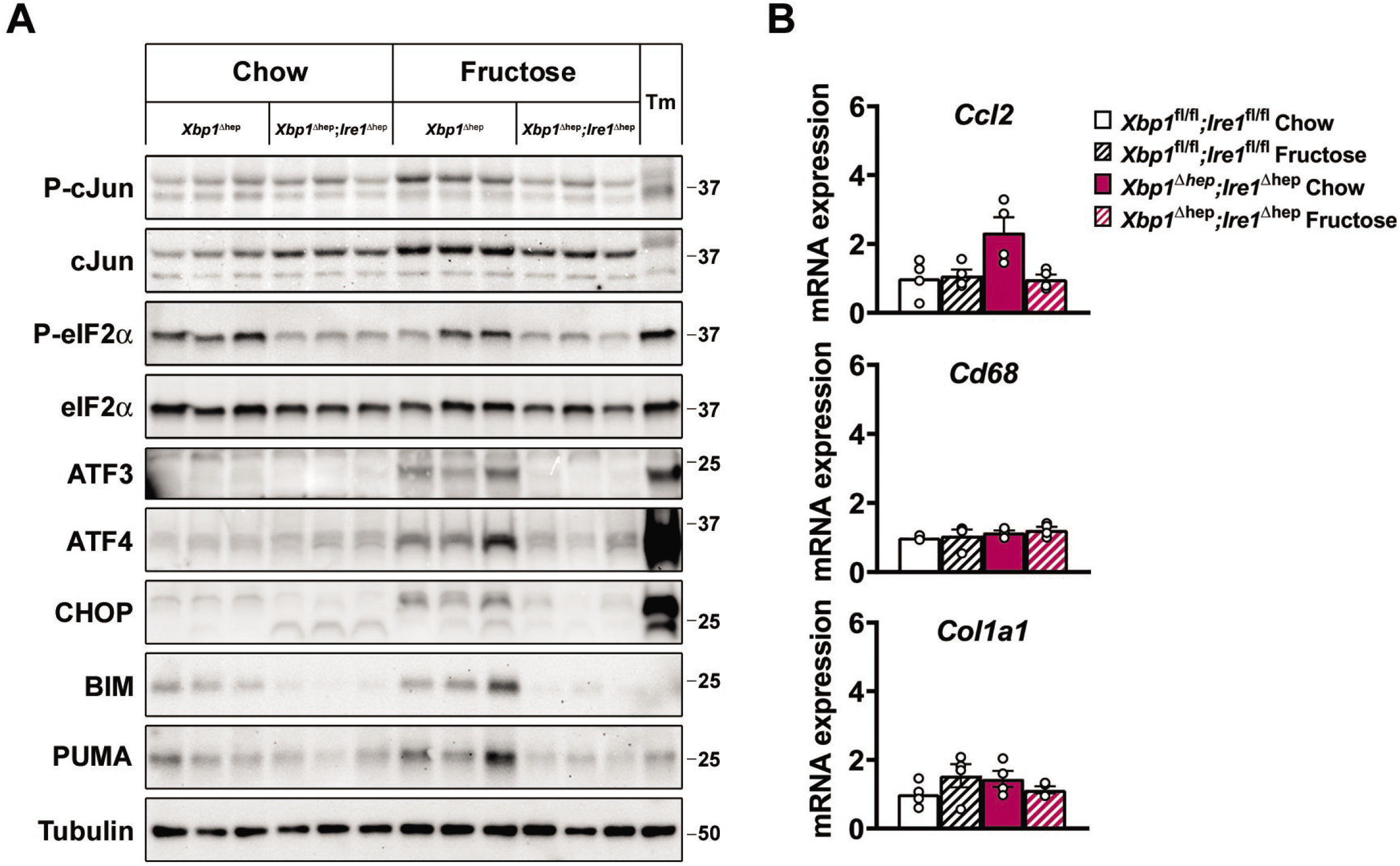
Comparison of ER stress responses in *Xbp1*^Δhep^ and *Xbp1*^Δhep^;*IRE1α*^Δhep^ mice in response to fructose feeding. **(A)** Western blots illustrate the expression of ER stress-related molecules in liver homogenates from *Xbp1*^Δhep^ and *Xbp1*^Δhep^;*IRE1α*^Δhep^ mice after 4 wk of chow or fructose feeding. Fructose-fed *Xbp1*^Δhep^ mice display phosphorylation of cJun and eIF2α as well as up-regulation of several eIF2α targets. These molecules are not induced in fructose-fed *Xbp1*^Δhep^;*IRE1α*^Δhep^ mice. **(B)** Graphs demonstrate the effect of fructose feeding for 4 wk on select genes pertinent to hepatic inflammation and fibrosis. Unlike *Xbp1*^Δhep^ mice, *Xbp1*^Δhep^;*IRE1α*^Δhep^ mice show no significant up-regulation of *Ccl2* mRNA following fructose feeding. There is also no induction of *Cd68* or *Col1a1*. Values represent mean ± SEM for n = 4.

To determine the long-term consequences of XBP1 deletion in hepatocytes, we challenged *Xbp1*^Δhep^ and *Xbp1*^Δhep^;*IRE1α*^Δhep^ mice with fructose vs. chow for 16 wk. Neither *Xbp1*^Δhep^ nor *Xbp1*^Δhep^;*IRE1α*^Δhep^ mice displayed any histologic signs of liver injury following 16 wk of chow feeding (**Figure 7A**). Fructose feeding for 16 wk caused modest hepatic steatosis in all 4 groups of mice; only in *Xbp1*^Δhep^ mice, however, was the steatosis associated with evidence of liver injury and early liver fibrosis (**Figure 7A, B**). *Xbp1*^Δhep^;*IRE1α*^Δhep^ mice were protected from the ALT elevations and fibrosis seen in *Xbp1*^Δhep^ mice (**Figure 7A, B**).

**Figure 7.**
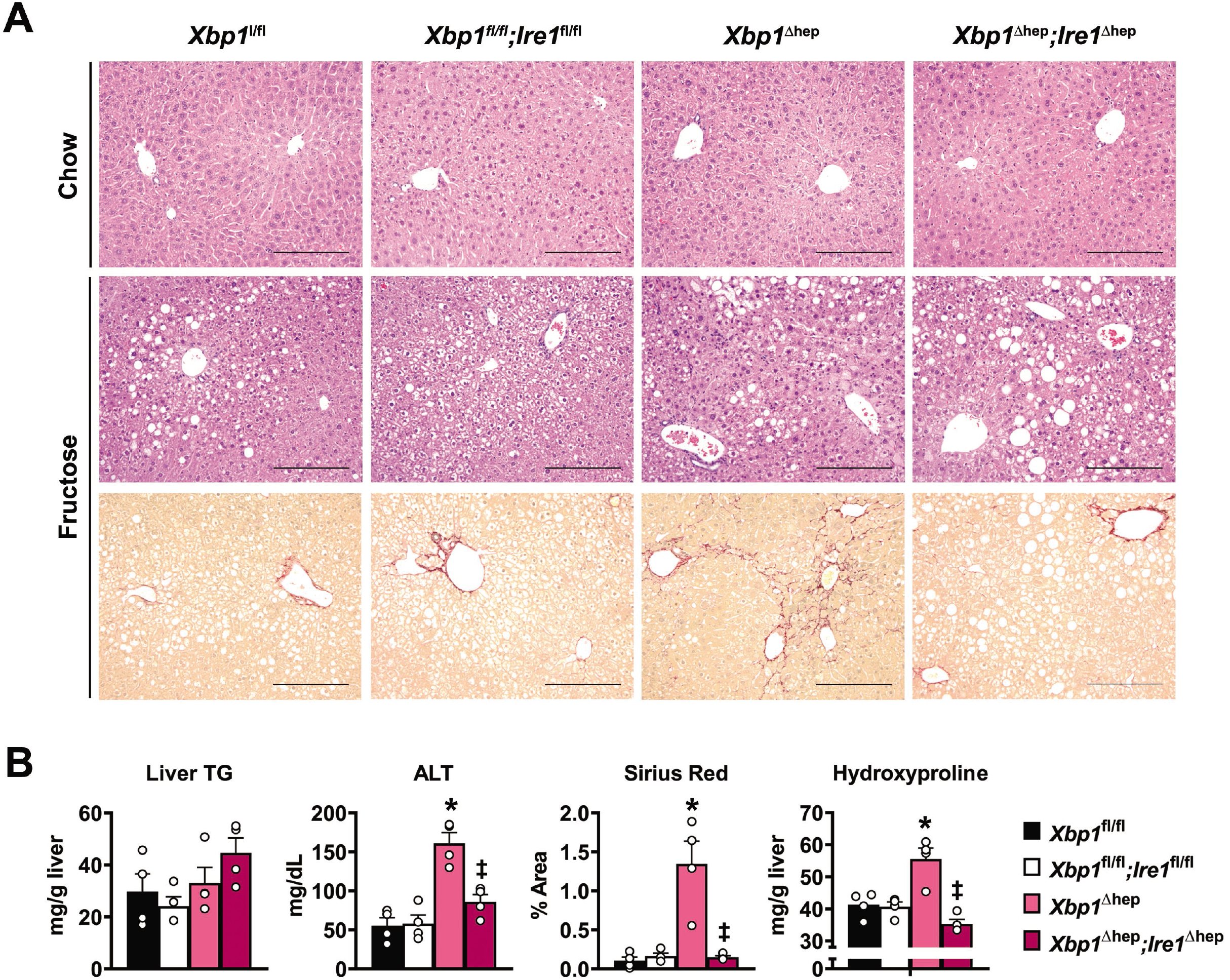
Long-term consequences of fructose feeding in *Xbp1*^Δhep^ and *Xbp1*^Δhep^;*IRE1α*^Δhep^ mice. **(A)** Photomicrographs demonstrate liver histology and fibrosis staining in *Xbp1*^Δhep^ and *Xbp1*^Δhep^;*IRE1α*^Δhep^ mice and their floxed controls after 16 wk of chow or fructose feeding. Chow-fed mice have no obvious histologic abnormalities; fructose feeding induced mild to moderate steatosis in all four groups of mice. Sirius red staining (bottom panels) show perisinusoidal and bridging fibrosis only in fructose-fed *Xbp1*^Δhep^ mice. Bar = 200 μm. **(B)** Graphs depict serum liver TG, serum ALT and fibrosis measured by morphometry and hydroxyproline content in control, *Xbp1*^Δhep^ and *Xbp1*^Δhep^;*IRE1α*^Δhep^ mice. The highest values of ALT and fibrosis are in fructose-fed *Xbp1*^Δhep^ mice at 16 wk. Values represent mean ± SEM for n = 4. *P* < 0.0005 by one-way ANOVA for ALT, Sirius Red morphometry and hydroxyproline. * *P* < 0.05 for *Xbp1*^Δhep^ vs. *Xbp1*^fl/fl^; ‡ *P* < 0.05 for *Xbp1*^Δhep^;*Ire1*^Δhep^ vs. *Xbp1*^Δhep^.

## Discussion

This study demonstrates that hepatocyte-specific deletion of XBP1 in adult mice renders them susceptible to acute and chronic liver injury. In *Xbp1*^Δhep^ mice, a mild dietary challenge that was insufficient to cause liver disease in control mice induced significant disease after gene deletion, characterized by hepatic steatosis, liver cell death and ultimately fibrosis. Importantly, the susceptibility of *Xbp1*^Δhep^ mice to liver disease was not the direct consequence of XBP1 deletion. Instead, it was due to compensatory up-regulation of the upstream ER stress transducer IRE1α. This was confirmed by liver-specific deletion of both XBP1 and IRE1α, which protected mice from the injury observed in mice lacking XBP1 alone.

Fructose-mediated liver injury in *Xbp1*^Δhep^ mice was associated with several features of heightened ER stress, including activation of IRE1α and eIF2α and the induction of the pro-apoptotic molecules JNK and CHOP. All of these stress responses were abrogated by IRE1α deletion, indicating they were triggered by IRE1α activation. Whether IRE1α provoked liver injury through its kinase or endoribonuclease functions or both is difficult to pinpoint: both were apparently up-regulated in our mice, based on the activation of JNK and RIDD. We suspect JNK was activated as a result of IRE1α kinase activity and eIF2α activated as a consequence of IRE1A endonuclease activity (RIDD) [23], although we acknowledge that these events could occur through other pathways [15,25,26]. We also cannot specify which of the several death-promoting molecules induced in *Xbp1*^Δhep^ livers were responsible for fructose-induced cell death. JNK, CHOP, BIM and PUMA were all up-regulated on a similar timeline, and thus they all likely contributed collectively to the development of liver injury.

One intriguing observation in the current study was that IRE1α and eIF2α were up-regulated at baseline in the livers of *Xbp1*^Δhep^ mice, yet there was no evidence of downstream signaling from these two molecules and no liver injury in the absence of an exogenous stimulus. This implies that some threshold of IRE1α activation must be surpassed in order to effect a cytotoxic response. Some have linked the cytotoxic potential of IRE1α to stress-induced high-order oligomerization of the protein with resultant activation of RIDD [27]; in the current study, though, RIDD was detectable in XBP1-deleted livers even before fructose feeding. RIDD-induced cell death may be cell-type specific [28,29], and hepatocytes may be somewhat resistant [11,30,31]. Still, our results support the concept that IRE1α can be pushed to a level that causes hepatotoxicity, and XBP1 deletion nearly achieves this goal.

Another relevant observation in our study was that IRE1α deletion did not protect *Xbp1*^Δhep^;*IRE1α*^Δhep^ mice from developing hepatic steatosis over 16 wk. This was likely due in part to the suppression of *Ph4b*, the gene encoding protein disulfide isomerase (PDI). PDI promotes the physiologic secretion of VLDL from the liver by controlling the activity of the microsomal triglyceride-transfer protein (MTP); downregulation of PDI predisposes to hepatic lipid accumulation [21]. Because *Ph4b* is a direct XBP1 target gene, its expression was low in both *Xbp1*^Δhep^ and *Xbp1*^Δhep^;*IRE1α*^Δhep^ mice (**see Figures 3B and S3C**). Still, the degree of liver injury was different in the two genetically engineered strains over the short and long term. This suggests that hepatic steatosis, although a feature of the liver injury seen in *Xbp1^Δhep^* mice, is not itself central to fructose-induced liver damage.

The current work uniquely supplements an existing body of research examining the impact of IRE1α and XBP1 on the liver. Previous studies of liver-targeted deletion of XBP1 have yielded mixed results: some showed no deleterious effect on the liver [8], others showed that XBP1 deletion improved hepatic steatosis in obese mice [11,32], and still others showed that XBP1 deletion worsened liver injury from pharmacologic or dietary insults [9,33]. Our experiments are in general agreement with the last group of reports indicating that XBP1 deletion is harmful to the liver when combined with another insult. However, our work diverges from these studies in that we attribute the harmful effect of hepatic XBP1 deletion to IRE1α. Regarding IRE1α, our work is in alignment with published studies demonstrating that its activity can protect against hepatic steatosis [21,34,35]. When IRE1α is activated to excess, however, it sensitizes hepatocytes to injurious stimuli.

In summary, the current work underscores the potential for IRE1α to induce liver injury when activated above a threshold level in the proper clinical setting. Indeed, IRE1α activity has been implicated as a contributing factor to lipotoxic liver injury and nonalcoholic fatty liver disease [36,37]. Still, not all research points to heightened IRE1α activity as a driving factor in liver disease pathogenesis; in some instances, S-nitrosylation of IRE1α, which impairs its endoribonuclease activity, has been linked to the development of hepatic steatosis [34,38]. Given the disconnection between hepatic steatosis and liver injury in our experiments, we conclude that IRE1α activation is detrimental to the liver independent of its effect on hepatic steatosis. IRE1α, therefore, may represent an important therapeutic target in hepatology.

## Acknowledgments

This work was supported in part by R01 DK068450 (JJM), T32 DK060414 (CCD), K08 DK098270 (ANM), a Pilot/Feasibility Award from the UCSF Liver Center (CCD) and an AASLD Pinnacle Award (CCD). The authors also acknowledge the support of the Cell Biology, Pathology and Immunology Cores of the UCSF Liver Center (P30 DK026743) and the Genome Core of the UCSF Helen Diller Family Comprehensive Cancer Center (P30 CA082103).

## Conflict of Interest

The authors declare no conflicts of interest.

---

**Figure S1. Phenotype of *Xbp1*^Δhep^ mice and response to tunicamycin**. **(A)** Western blots demonstrate the responses of *Xbp1*^fl/fl^ and *Xbp1*^Δhep^ mice to a single dose of tunicamycin (Tm) in vivo (1 mg/kg x 3 h). XBP1s and lamin were measured in liver nuclear extracts; the remaining proteins were measured in whole liver homogenates. Nuclear translocation of XBP1s was prominent in *Xbp1*^fl/fl^ livers but nearly absent in *Xbp1*^Δhep^ livers and completely absent in an untreated control (C). Tunicamycin-induced phosphorylation of IRE1α and cJun was not impaired in *Xbp1*^Δhep^ livers. **(B)** Graph confirms that that tunicamycin induces XBP mRNA splicing in *Xbp1*^fl/fl^ and *Xbp1*^Δhep^ livers. In *Xbp1*^Δhep^ livers, the Cre-mediated gene deletion does not involve the mRNA splice site [8]; thus, mRNA splicing can occur but XBP1s protein is not effectively translated. **(C)** Graphs demonstrate reduced serum lipids in *Xbp1*^Δhep^ mice compared to *Xbp1*^fl/fl^ mice under basal conditions. **(D)** Photomicrographs illustrate normal liver histology in *Xbp1*^fl/fl^ and *Xbp1*^Δhep^ mice under basal conditions. Bar = 200 μm. Values represent mean ± SEM. * *P* < 0.05 for Tm vs control; ‡ *P* < 0.05 for *Xbp1*^Δhep^ vs. *Xbp1*^fl/fl^.

**Figure S2. Lipogenic gene expression after fructose feeding for 4 weeks**. Graph depicts lipogenic gene expression in the livers of *Xbp1*^fl/fl^ and *Xbp1*^Δhep^ mice after 4 wk of fructose feeding. Values represent mean ± SEM for n = 4. * *P* < 0.05 for fructose vs. chow of same genotype; ‡ *P* < 0.05 for *Xbp1*^Δhep^ vs. *Xbp1*^fl/fl^.

**Figure S3. Verification of IRE1α deletion and the absence of XBP1 splicing or RIDD in *Xbp1*^Δhep^;*IRE1α*^Δhep^ livers**. **(A)** Western blot illustrates the presence of a truncated form of IRE1α in liver homogenates from *Xbp1*^Δhep^;*IRE1α*^Δhep^ mice, representing IRE1α lacking the RNase domain [16,24]. **(B)** Graph depicts a lack of spontaneous Xbp1 mRNA splicing in *Xbp1*^Δhep^;*IRE1α*^Δhep^ livers, in contrast to *Xbp1*^Δhep^ livers (see figure S1B). (C) Graphs demonstrate that direct XBP1 target genes are suppressed as expected in *Xbp1*^Δhep^;*IRE1α*^Δhep^ livers, but RIDD target genes are not suppressed due to the concomitant absence of the IRE1α endoribonuclase. Values represent mean ± SEM for n = 4. By unpaired t-test, * *P* < 0.05 for *Xbp1*^Δhep^;*IRE1α*^Δhep^ vs. *Xbp1*^fl/fl^;*IRE1α*^fl/fl^.

## References

1 Lee, AH, Iwakoshi, NNGlimcher, LH. XBP-1 regulates a subset of endoplasmic reticulum resident chaperone genes in the unfolded protein response. Mol Cell Biol 2003; 23: 7448–7459.

2 Sriburi, R, Jackowski, S, Mori, KBrewer, JW. XBP1: a link between the unfolded protein response, lipid biosynthesis, and biogenesis of the endoplasmic reticulum. J Cell Biol 2004; 167: 35–41.

3 Sriburi, R, Bommiasamy, H, Buldak, GL, Robbins, GR, Frank, M, Jackowski, S et al. Coordinate regulation of phospholipid biosynthesis and secretory pathway gene expression in XBP-1(S)-induced endoplasmic reticulum biogenesis. J Biol Chem 2007; 282: 7024–7034.

4 Reimold, AM, Iwakoshi, NN, Manis, J, Vallabhajosyula, P, Szomolanyi-Tsuda, E, Gravallese, EM et al. Plasma cell differentiation requires the transcription factor XBP-1. Nature 2001; 412: 300–307.

5 Kaser, A, Lee, AH, Franke, A, Glickman, JN, Zeissig, S, Tilg, H et al. XBP1 links ER stress to intestinal inflammation and confers genetic risk for human inflammatory bowel disease. Cell 2008; 134: 743–756.

6 Ozcan, L, Ergin, AS, Lu, A, Chung, J, Sarkar, S, Nie, D et al. Endoplasmic reticulum stress plays a central role in development of leptin resistance. Cell Metab 2009; 9: 35–51.

7 Jurczak, MJ, Lee, AH, Jornayvaz, FR, Lee, HY, Birkenfeld, AL, Guigni, BA et al. Dissociation of inositol-requiring enzyme (IRE1alpha)-mediated c-Jun N-terminal kinase activation from hepatic insulin resistance in conditional X-box-binding protein-1 (XBP1) knock-out mice. J Biol Chem 2012; 287: 2558–2567.

8 Lee, AH, Scapa, EF, Cohen, DEGlimcher, LH. Regulation of hepatic lipogenesis by the transcription factor XBP1. Science 2008; 320: 1492–1496.

9 Olivares, SHenkel, AS. Hepatic Xbp1 Gene Deletion Promotes Endoplasmic Reticulum Stress-induced Liver Injury and Apoptosis. J Biol Chem 2015; 290: 30142–30151.

10 Argemi, J, Kress, TR, Chang, HCY, Ferrero, R, Bertolo, C, Moreno, H et al. X-box Binding Protein 1 Regulates Unfolded Protein, Acute-Phase, and DNA Damage Responses During Regeneration of Mouse Liver. Gastroenterology 2017; 152: 1203–1216 e1215.

11 So, JS, Hur, KY, Tarrio, M, Ruda, V, Frank-Kamenetsky, M, Fitzgerald, K et al. Silencing of lipid metabolism genes through IRE1alpha-mediated mRNA decay lowers plasma lipids in mice. Cell Metab 2012; 16: 487–499.

12 Hetz, C, Chevet, EOakes, SA. Proteostasis control by the unfolded protein response. Nat Cell Biol 2015; 17: 829–838.

13 Rutkowski, DT, Wu, J, Back, SH, Callaghan, MU, Ferris, SP, Iqbal, J et al. UPR pathways combine to prevent hepatic steatosis caused by ER stress-mediated suppression of transcriptional master regulators. Dev Cell 2008; 15: 829–840.

14 Hollien, J, Lin, JH, Li, H, Stevens, N, Walter, PWeissman, JS. Regulated Ire1-dependent decay of messenger RNAs in mammalian cells. J Cell Biol 2009; 186: 323–331.

15 Tabas, IRon, D. Integrating the mechanisms of apoptosis induced by endoplasmic reticulum stress. Nat Cell Biol 2011; 13: 184–190.

16 Iwawaki, T, Akai, R, Yamanaka, SKohno, K. Function of IRE1 alpha in the placenta is essential for placental development and embryonic viability. Proc Natl Acad Sci U S A 2009; 106: 16657–16662.

17 Pickens, MK, Yan, JS, Ng, RK, Ogata, H, Grenert, JP, Beysen, C et al. Dietary sucrose is essential to the development of liver injury in the methionine-choline-deficient model of steatohepatitis. J Lipid Res 2009; 50: 2072–2082.

18 Folch, J, Lees, MSloane Stanley, GH. A simple method for the isolation and purification of total lipides from animal tissues. J Biol Chem 1957; 226: 497–509.

19 Lee, GS, Yan, JS, Ng, RK, Kakar, SMaher, JJ. Polyunsaturated fat in the methionine-choline-deficient diet influences hepatic inflammation but not hepatocellular injury. J Lipid Res 2007; 48: 1885–1896.

20 Jamall, IS, Finelli, VNQue Hee, SS. A simple method to determine nanogram levels of 4-hydroxyproline in biological tissues. Anal Biochem 1981; 112: 70–75.

21 Wang, S, Chen, Z, Lam, V, Han, J, Hassler, J, Finck, BN et al. IRE1alpha-XBP1s induces PDI expression to increase MTP activity for hepatic VLDL assembly and lipid homeostasis. Cell Metab 2012; 16: 473–486.

22 Hetz, C. The unfolded protein response: controlling cell fate decisions under ER stress and beyond. Nat Rev Mol Cell Biol 2012; 13: 89–102.

23 So, JS, Cho, S, Min, SH, Kimball, SRLee, AH. IRE1alpha-Dependent Decay of CReP/Ppp1r15b mRNA Increases Eukaryotic Initiation Factor 2alpha Phosphorylation and Suppresses Protein Synthesis. Mol Cell Biol 2015; 35: 2761–2770.

24 Tsuchiya, Y, Saito, M, Kadokura, H, Miyazaki, JI, Tashiro, F, Imagawa, Y et al. IRE1-XBP1 pathway regulates oxidative proinsulin folding in pancreatic beta cells. J Cell Biol 2018; 217: 1287–1301.

25 Taniuchi, S, Miyake, M, Tsugawa, K, Oyadomari, MOyadomari, S. Integrated stress response of vertebrates is regulated by four eIF2alpha kinases. Sci Rep 2016; 6: 32886.

26 Timmins, JM, Ozcan, L, Seimon, TA, Li, G, Malagelada, C, Backs, J et al. Calcium/calmodulin-dependent protein kinase II links ER stress with Fas and mitochondrial apoptosis pathways. J Clin Invest 2009; 119: 2925–2941.

27 Ghosh, R, Wang, L, Wang, ES, Perera, BG, Igbaria, A, Morita, S et al. Allosteric inhibition of the IRE1alpha RNase preserves cell viability and function during endoplasmic reticulum stress. Cell 2014; 158: 534–548.

28 Lerner, AG, Upton, JP, Praveen, PV, Ghosh, R, Nakagawa, Y, Igbaria, A et al. IRE1alpha induces thioredoxin-interacting protein to activate the NLRP3 inflammasome and promote programmed cell death under irremediable ER stress. Cell Metab 2012; 16: 250–264.

29 Upton, JP, Wang, L, Han, D, Wang, ES, Huskey, NE, Lim, L et al. IRE1alpha cleaves select microRNAs during ER stress to derepress translation of proapoptotic Caspase-2. Science 2012; 338: 818–822.

30 Hur, KY, So, JS, Ruda, V, Frank-Kamenetsky, M, Fitzgerald, K, Koteliansky, V et al. IRE1alpha activation protects mice against acetaminophen-induced hepatotoxicity. J Exp Med 2012; 209: 307–318.

31 Maurel, M, Chevet, E, Tavernier, JGerlo, S. Getting RIDD of RNA: IRE1 in cell fate regulation. Trends Biochem Sci 2014; 39: 245–254.

32 Herrema, H, Zhou, Y, Zhang, D, Lee, J, Salazar Hernandez, MA, Shulman, GI et al. XBP1s Is an Anti-lipogenic Protein. J Biol Chem 2016; 291: 17394–17404.

33 Liu, X, Henkel, AS, LeCuyer, BE, Schipma, MJ, Anderson, KAGreen, RM. Hepatocyte X-box binding protein 1 deficiency increases liver injury in mice fed a high-fat/sugar diet. Am J Physiol Gastrointest Liver Physiol 2015; 309: G965–974.

34 Wang, JM, Qiu, Y, Yang, Z, Kim, H, Qian, Q, Sun, Q et al. IRE1alpha prevents hepatic steatosis by processing and promoting the degradation of select microRNAs. Sci Signal 2018; 11.

35 Zhang, K, Wang, S, Malhotra, J, Hassler, JR, Back, SH, Wang, G et al. The unfolded protein response transducer IRE1alpha prevents ER stress-induced hepatic steatosis. EMBO J 2011; 30: 1357–1375.

36 Kakazu, E, Mauer, AS, Yin, MMalhi, H. Hepatocytes release ceramide-enriched pro-inflammatory extracellular vesicles in an IRE1alpha-dependent manner. J Lipid Res 2016; 57: 233–245.

37 Lebeaupin, C, Vallee, D, Rousseau, D, Patouraux, S, Bonnafous, S, Adam, G et al. Bax inhibitor-1 protects from nonalcoholic steatohepatitis by limiting inositol-requiring enzyme 1 alpha signaling in mice. Hepatology 2018; 68: 515–532.

38 Yang, L, Calay, ES, Fan, J, Arduini, A, Kunz, RC, Gygi, SP et al. METABOLISM. S-Nitrosylation links obesity-associated inflammation to endoplasmic reticulum dysfunction. Science 2015; 349: 500–506.

